# Modelling the tumor metallome *in vitro* reveals manganese as a unique tumor-promoting element

**DOI:** 10.1101/2025.09.15.676369

**Authors:** Isabella C. Abrahão dos Santos, Carolina Araújo Pereira da Silva, Déborah Assis de A. Carneiro, Arthur Moreira da Rocha, Carlos Pérez, Simone C. Cardoso, Mariana P. Stelling

## Abstract

The microenvironment (TME) undergoes significant modification in tumor progression. TME comprises locally secreted molecules, cells, and metal ions. Metals interact with the cell surface or are internalized, modulating downstream pathways related to adhesion, migration, and proliferation. Our group showed that tumor-bearing mice present altered Mn distribution. Mn-exposed tumor cells showed *in vitro* higher proliferation and migration, associated with changes in glycocalyx organization. In this work, we modulated, *in vitro*, tumor cells’ metallome, and evaluated metals internalization, cellular distribution and its early effect on migration. Cell viability and survival were evaluated using the MTT and clonogenic assays. Metal retention and distribution were assessed with inductively coupled plasma optical emission spectroscopy (ICP-OES) and high-resolution X-ray fluorescence (XRF). Cell migration was evaluated with wound healing assay. Our results indicate that an early interaction between Mn and the cell surface, probably the negatively-charged glycocalyx, induces morphological changes that lead to increased invasiveness. Future investigations will help to better understand the mechanisms of Mn retention and internalization during tumor progression.

## Introduction

**T**he tumor microenvironment (TME) is deeply disturbed during tumor progression as a consequence of genetic and epigenetic alterations. The cellular population in the TME is highly heterogeneous, comprising tumor cells, cancer stem cells (CSCs), healthy cells, and immune infiltrating cells, which alters the gradient of signaling molecules and results in a favorable environment for tumor progression^1^. This disturbance in TME results in altered expression of receptors and transporters on the cell surface, which are strongly related to cell invasion and metastasis formation^2,3^, for example. Among these changes, an increase in the availability of nutrients, such as glucose, soluble proteins, and trace elements, is also observed ^4–6^. Trace elements such as zinc, iron, and magnesium are considered micronutrients due to their nutritional significance and essentiality. These elements are essential for cellular roles as (1) enzymatic cofactors; (2) protein stabilization; (3) DNA replication and repair; (4) reactive oxygen species (ROS) neutralization, among others^7–9^.

Changes in the concentration and distribution of trace elements have been linked to cancer. Eating habits, smoking, and environmental exposure are the main causes associated with the accumulation of metals, such as copper and zinc, with the development of cancer, as well as tumor accumulation of these elements^10^. Elements as manganese, iron, and copper were found significantly elevated in prostate, lung, pancreatic, esophageal, liver, and breast cancer samples when compared to their healthy counterparts^11–15^. The use of sample microarrays is also interesting for understanding the spatial distribution and accumulation of metals in tumor and adjacent tissues using spectroscopic techniques^16^. With this approach, Doble & Miklos observed that radioresistant tumors like glioblastoma and melanoma presented the highest concentrations of manganese, and were associated with the worst survival rate^17^.

In addition to tissue changes, recent research has been investigating alterations in the metallome of biological fluids for cancer screening. Serum and urine from patients with gallbladder, thyroid, lung, breast, prostate, and pancreatic cancer presented a specific metallome profile when compared to healthy individuals^14,18–23^, suggesting a new non-invasive method for screening tumor progression or organ failure due to cancer, especially if associated with other metabolite and enzyme analysis^20,24,25^.

*In vitro* and *in silico* studies are also evaluating the status of genes, transcripts, and proteins related to metal metabolism. Transcriptomic analysis of breast and prostate cancer, as well as myeloid leukemia cell lineages, revealed alterations in zinc (Zn) metabolism, characterized by an increase in ZIP (ZRT, IRT-like protein) transporters and metallothionein (MTs), accompanied by a concomitant reduction in ZnT (*Zn transporter*)^26–28^. Patient data obtained from The Cancer Genome Atlas (TCGA) revealed an altered pattern of genes related to manganese metabolism and immune system regulation, which can stratify patients with gastric cancer into high- and low-risk groups based on gene expression and also predict their response to chemotherapy and immunotherapy. TCGA analysis also indicated the transcriptional factor (TF) copper-dependent ATOX1 as a potential cause of copper intracellular accumulation in breast cancer cells in triple-negative (TNBC) and HER2 subtypes^29^ and proliferative activity in non-Hodgkin tumor cells^30^.

The use of tumor xenografts in animal models is also extremely important to understand elemental distribution and how this contributes to tumor progression, and even if the metallomic profile of distant tissues is altered. Both studies by Planeta et al. with rats suggest that metallomic changes in brain tissues are correlated with glioblastoma (GBM) growth and invasiveness, where the tumor area presents an increase in iron and selenium availability while the surrounding area exhibits copper accumulation^31,32^. Previously, our group demonstrated that metastatic Lewis’ lung carcinoma modulates manganese distribution in primary tumor and metastatic sites as the liver and lungs^33^. Later, the elemental profile of distant tissues was evaluated. We reported that the primary and distant sites exhibited a noticeable increase in iron and manganese content over the weeks. Additionally, a discrete increase in copper concentration was observed, while zinc showed a stable distribution with no statistical difference^34^. In this work, we aim to evaluate the effect of metallomic imbalance on tumor cells’ behavior by simulating these events *in vitro*. We expect to evaluate cells’ sensitivity to high concentrations of manganese, copper, iron, and zinc, as well as investigate intracellular accumulation and distribution of these elements, and finally understand if these elements can promote an invasive behavior.

## Results and discussion

### 1. Tumor cell metabolism and survival in the presence of increasing concentrations of MnCl_2_, FeCl_2_, CuCl_2_, and ZnCl_2_

To further understand the effects of metal ion accumulation in lung cancer cells, we first evaluated cell metabolism and survival after 24h exposure to increasing concentrations of MnCl_2_, FeCl2, CuCl2, or ZnCl2. For metabolism evaluation, we performed the MTT assay 24h after cell exposure from 1.5 to 500 µM of salts added to the culture medium. Cells exposed to concentrations above 12.5 µM of MnCl2 exhibited high sensitivity to Mn toxicity, with higher concentrations resulting in a significant decrease in viability compared to the control group (Figure 1). Interestingly, only the concentration of 6.2 µM MnCl_2_ did not affect cell metabolism, while 1.5 µM and 3.5 µM reduced viability significantly by 39.1% and 25.5%, respectively (Figure 1). When cultivated in the presence of CuCl_2_, LLC cells exhibited reduced viability at every concentration tested, except at 12.5, 25, 50, and 100 µM. However, viability decreased to less than 50% only at 500 µM (Figure 1). Cells exposed to FeCl_2_ exhibited an interesting behavior: lower concentrations (1.5 to 6.2 µM) impaired cell viability, whereas concentrations above 12.5 µM (except 500 µM) did not affect cell viability (Figure 1). Finally, cells cultivated with ZnCl_2_ supplementation only had decreased viability when exposed to 500 µM (Figure 1).

**Figure 1.**
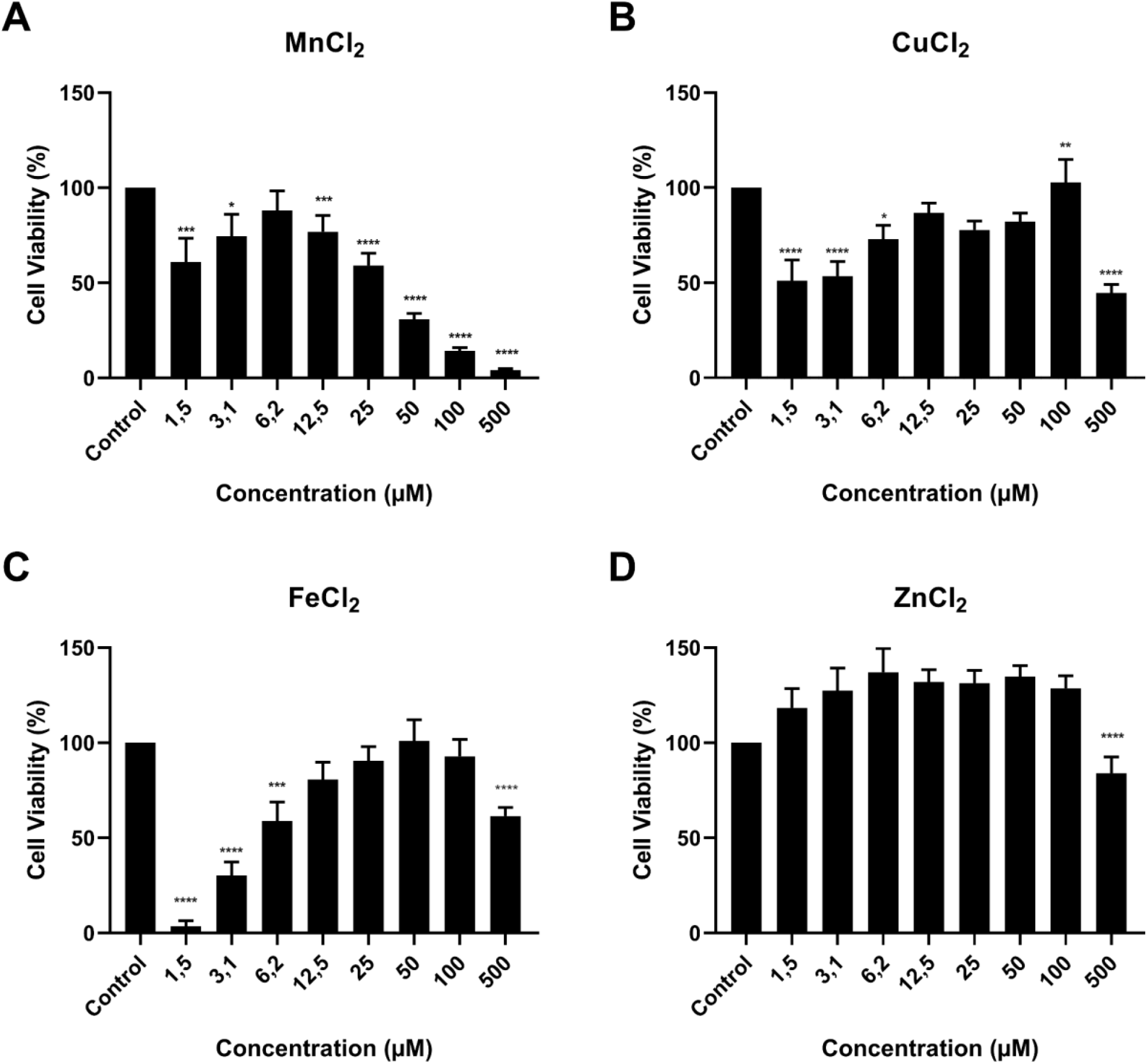
Quantification of cell viability using the MTT assay. After 24 hours of exposure to serial concentrations of (A) MnCl_2_, (B) CuCl_2_, (C) FeCl_2_, and (D) ZnCl_2_, cell viability was quantified using the colorimetric reaction of the MTT assay. * p≤0.05; ** p≤0.01; *** p≤0.001; **** p ≤0.0.0001. Mean ± Standard Error. n=5. One-Way ANOVA with Dunnett’s post-test.

Next, we performed the clonogenic assay to confirm whether these changes in cell metabolism caused by metallomic imbalance could influence cell survival. Cells were cultivated for 24h in regular medium (DMEM high glucose) or supplemented with 1, 5, 50, or 500 µM of MnCl_2_, FeCl_2_, CuCl_2_, or ZnCl_2_. Cells cultivated in 1 or 5 µM of MnCl_2_ maintained 93.5% and 96.7%, respectively, of survival, while the groups exposed to 50 and 500 µM had 53.4% and 0.4% survival compared to the control group (Figure 2, B). Cells exposed to FeCl_2_ maintained survival above 90% with all concentrations tested (Figure 2, C). For the cells cultivated with CuCl_2_, the groups cultivated in 50 and 500 µM decreased by 14.5% and 49.2% (Figure 2, D). Except for the 500 µM concentration, all cells exposed to ZnCl_2_ presented morethan 90% survival (Figure 2, E). Based on MTT and clonogenic data, we performed all the experiments with 5 µM of the salts.

**Figure 2.**
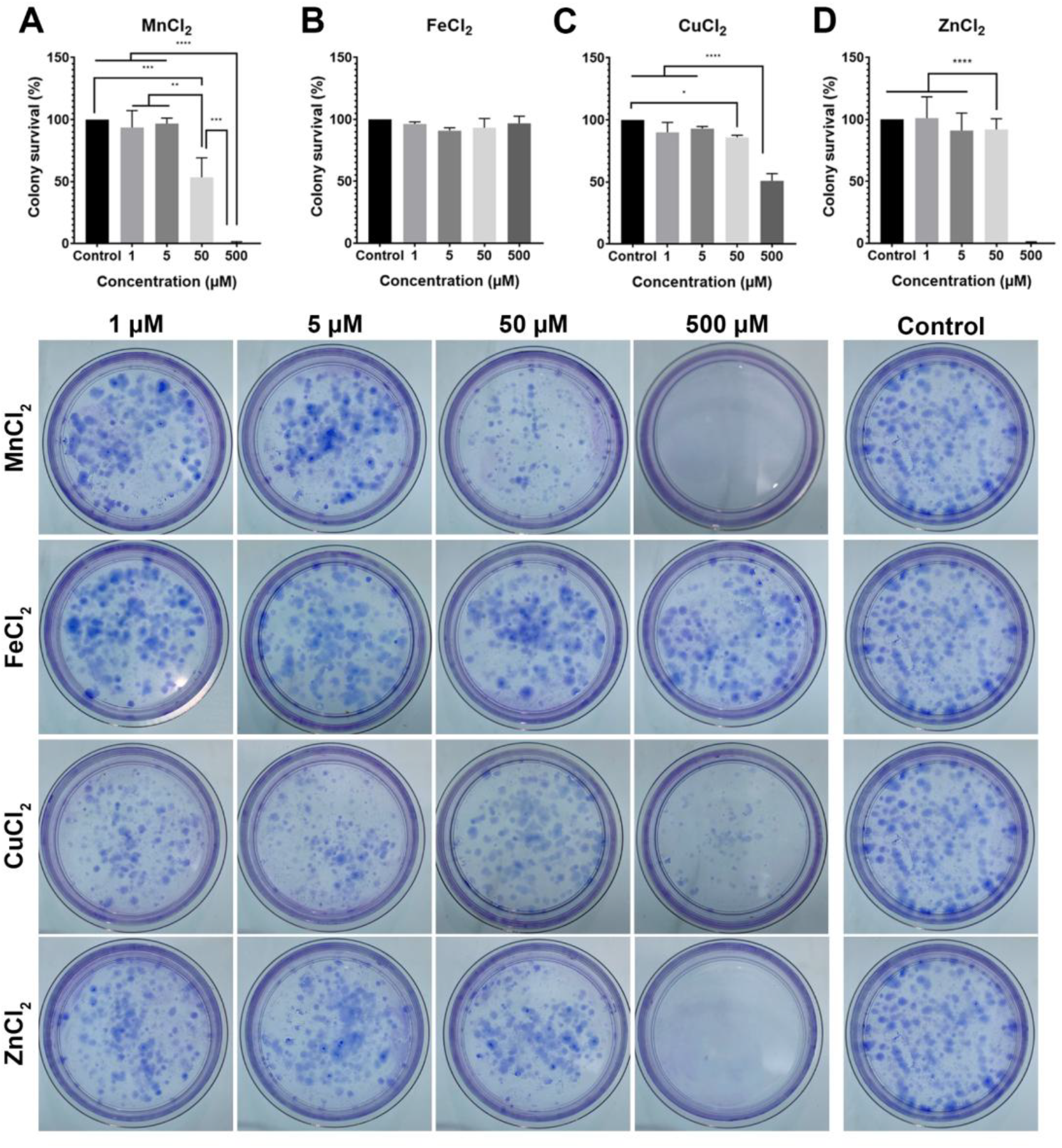
Cell survival using the clonogenic assay after 24 hours of exposure to 1, 5, 50, or 500 µM of (A) MnCl_2_, (B) FeCl_2_, (C) CuCl_2_, and (D) ZnCl_2_. Representative images of crystal violet-stained colonies from the control groups or those exposed to different salt concentrations. **p≤0.01; ***p≤0.001; **** p≤0.0.0001. Mean ± standard deviation. n=3. One-way ANOVA with Tukey’s post-test.

Results obtained for Mn crescent concentrations for both MTT and clonogenic assays were according with previous data obtained by our group, where concentrations between 5 and 12.5 µM were safe for cells. Concentrations above 25 µM significantly reduced cell viability and survival^33^. These data also suggest that elevated Mn concentrations are toxic to tissues other than neurological^34,35^. Recent evidence suggests that Mn toxicity occurs through ferroptosis, and this mechanism is being investigated to develop new antitumoral strategies.

We also observed a pattern of resistance to toxicity mediated by high concentrations of Fe and Cu (100 and 500 µM), suggesting an evasion of ferroptosis and cuproptosis. This was previously observed in cellular models of lung^36^, pancreatic^37^, endometrial^38^, and colorectal^39^cancer. This phenomenon has aroused interest in research for prognostic biomarkers related to ferroptosis and ferroptosis resistance^40^, as well as the development of therapies to reverse ferroptosis resistance in cancer^41^. In non-resistant models, the use of Cu ionophores, as well as nanoparticles to overload tumor cells with Fe and Cu, and the discovery of ferroptosis inducer compounds, have been studied to complement traditional therapy in hepatocellular carcinoma^42^, non-small-cell lung carcinoma^43^, neuroblastoma^44^, glioma^45^, colon adenocarcinoma, and TNBC^46^ models.

For Zn, both the MTT and clonogenic assays exhibited high viability and survival up to 50 µM. Regardless, our data for 500 µM were contradictory between the MTT and clonogenic assays (Figures 1D and 2D) after 24-hour exposure. Commonly, Zn is associated with a protective role for cells, as this element is a cofactor for several enzymes related to redox activity. However, this element is also associated with neuron death via mitochondrial damage^47–50^. Zn overload causes ROS generation and glycolysis inhibition, resulting in decreased NAD^+^ levels and reduction of the tricarboxylic acid cycle (TCA) ^50,51^. An attenuated event was observed in astrocytes^52^. Nevertheless, in vitro data for Zn toxicity, and even other metals, may not be completely reliable due to cell culture conditions, as the presence of serum, which contains cation-binding proteins such as albumine^53,54^.

Herein, data presented for viability and survival suggests how LLC cells respond to extreme metallome alterations. Further experiments with chronic exposure to these metals may elicit tumor tolerance to alterations in the TME metallome in the long term.

### 2. LLC cells rapidly internalize Mn ions, which present a heterogeneous distribution

Once the concentration of the salts was defined, we subjected cells to 1 or 4h exposure to the metals at 5 µM, and quantified the element rate through ICP-OES, to understand whether LLC cells are capable of internalizing and accumulating metals. Notably, cells cultivated with MnCl_2_presented the highest ratio of intracellular accumulation compared to the control group (Figure 3, A). Table 2 demonstrates that the addition of salt to the medium did not significantly interfere with the concentration of other elements (e.g., FeCl_2_ addition did not alter the intracellular concentrations of Mn or Zn).

**Figure 3.**
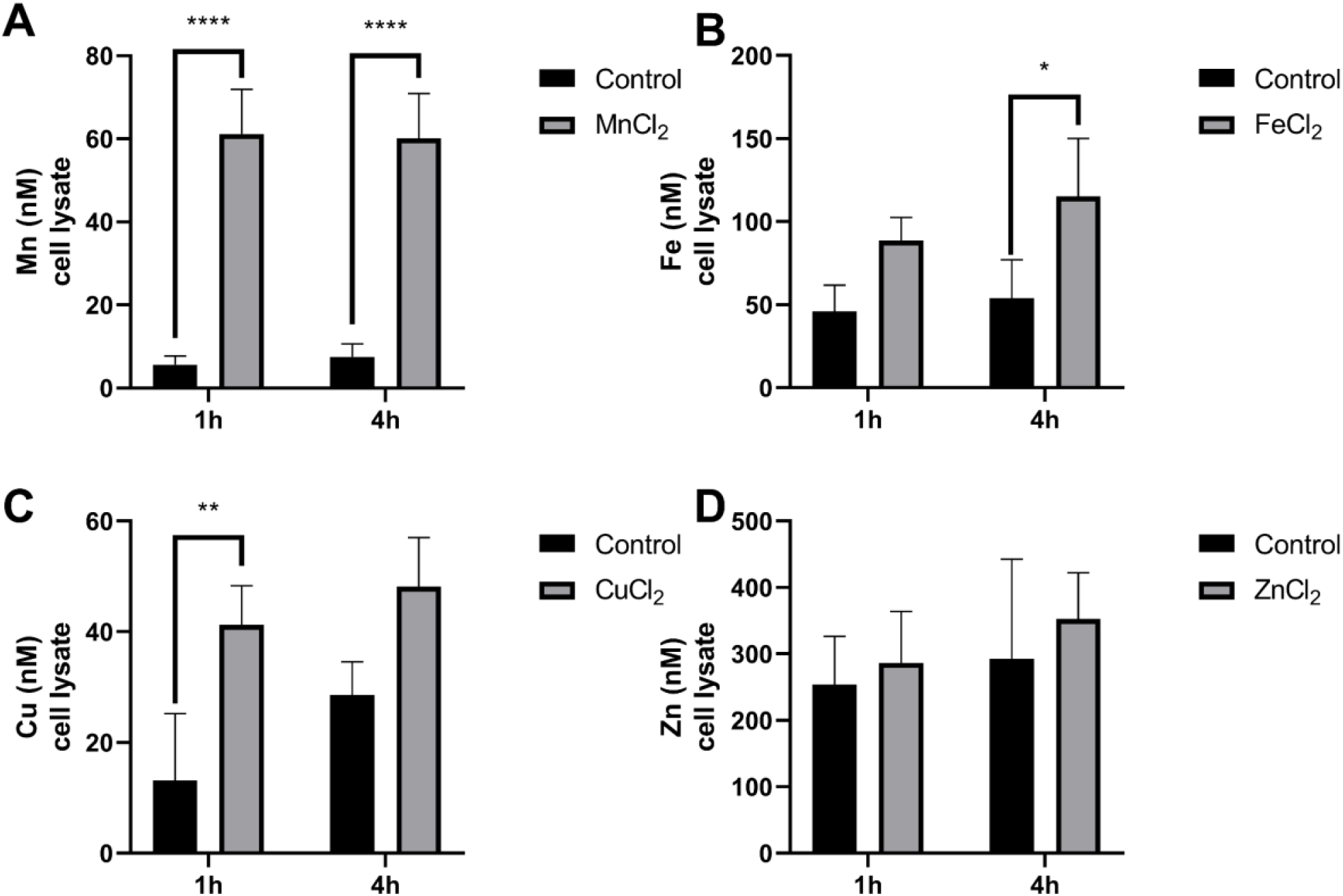
Metal content quantification in the cell lysate after 1 and 4h exposure to 5 µM of (A) MnCl_2_, (B) FeCl_2_, (C) CuCl_2_, or (D) ZnCl_2_. *p≤0.05; **p≤0.01; **** p≤0.0.0001. Mean ± standard deviation. n=4. One-way ANOVA with Sidak’s multiple comparison post-test.

|  | **Zn (nM)** | **Mn (nM)** | **Fe (nM)** | **Cu (nM)** | **K (mM)** | **Na (mM)** |
| --- | --- | --- | --- | --- | --- | --- |
| **Control -1h** | 253.9 | 5.64 | 46.08 | 13.12 | 0.12 | 0.51 |
| **ZnCl<sub>2</sub> - 1h** | 286.33 | 7.52 | 55.99 | 20.41 | 0.13 | 0.75 |
| **MnCl<sub>2</sub> - 1h** | 229.63 | 61.16 | 34.86 | 16.84 | 0.12 | 0.45 |
| **FeCl<sub>2</sub> -1h** | 263.18 | 10.19 | 88.59 | 18.83 | 0.13 | 0.65 |
| **CuCl<sub>2</sub> - 1h** | 301.11 | 7.46 | 52.65 | 41.23 | 0.14 | 0.76 |
| **Control - 4h** | 292.9 | 7.52 | 53.96 | 28.59 | 0.14 | 0.49 |
| **ZnCl<sub>2</sub> -4h** | 352.66 | 5.04 | 74.14 | 33.36 | 0.10 | 0.61 |
| **MnCl<sub>2</sub> - 4h** | 311.21 | 60.14 | 49.25 | 27.75 | 0.09 | 0.41 |
| **FeCl<sub>2</sub> - 4h** | 306.77 | 12.13 | 115.27 | 29.9 | 0.10 | 0.03 |
| **CuCl<sub>2</sub> - 4h** | 321.56 | 5.4 | 64.47 | 48.11 | 0.09 | 0.05 |

Cells cultivated during 1h in MnCl_2_ presented 11X and 4h 8.5X more manganese than cells grown in basal medium (Figure 3, A). Mn concentration in the control group was 5.6 and 7.5 nM, and after 1h and 4h, exposure increased to 61.1 and 60.1 nM, respectively (Figure 3, A). While cells cultivated upon FeCl_2_, CuCl_2_, or ZnCl_2_ increased from 1.5 to 2X of their respective element (Figure 3, B, D, and D). The FeCl_2_ group presented a higher increase in Fe content after 4h, with 115.2 nM, while cells exposed for 1h had 88.5 nM, and control samples varied from 46 nM and 53.9 nM, respectively (Figure 3, B). Cells exposed to CuCl_2_ for 1h increased Cu concentration from 13.1 nM (control) to 41.4 nM (Figure 3, C). Finally, cells cultivated in media supplemented with ZnCl_2_ did not present any significant change in Zn concentration (Figure 3, D).

Once we detected that only the MnCl_2_ group presents a high rate of internalization, we next investigated elemental distribution in our samples through high-resolution X-ray fluorescence (XRF). Cells were grown in control medium or medium supplemented with MnCl_2_ 5 µM for 1h or 4h and then irradiated at Carnaúba beamline (LNLS). First, we obtained the multielemental maps in ‘panorama’ mode to obtain a broad view of our samples (Figure 4). The panorama map is composed of 16 regions of 50×50 µm irradiated and processed to generate a 200×200 µm map, with 800 nm resolution. In these maps, we observed cell aggregates delimited by the element K, in red. Comparing the three conditions, we observed that the control group showed few niches rich in Mn (green) or Fe (blue), while both time points of the MnCl_2_ group showed an increase in Mn-rich niches (Figure 4).

**Figure 4.**
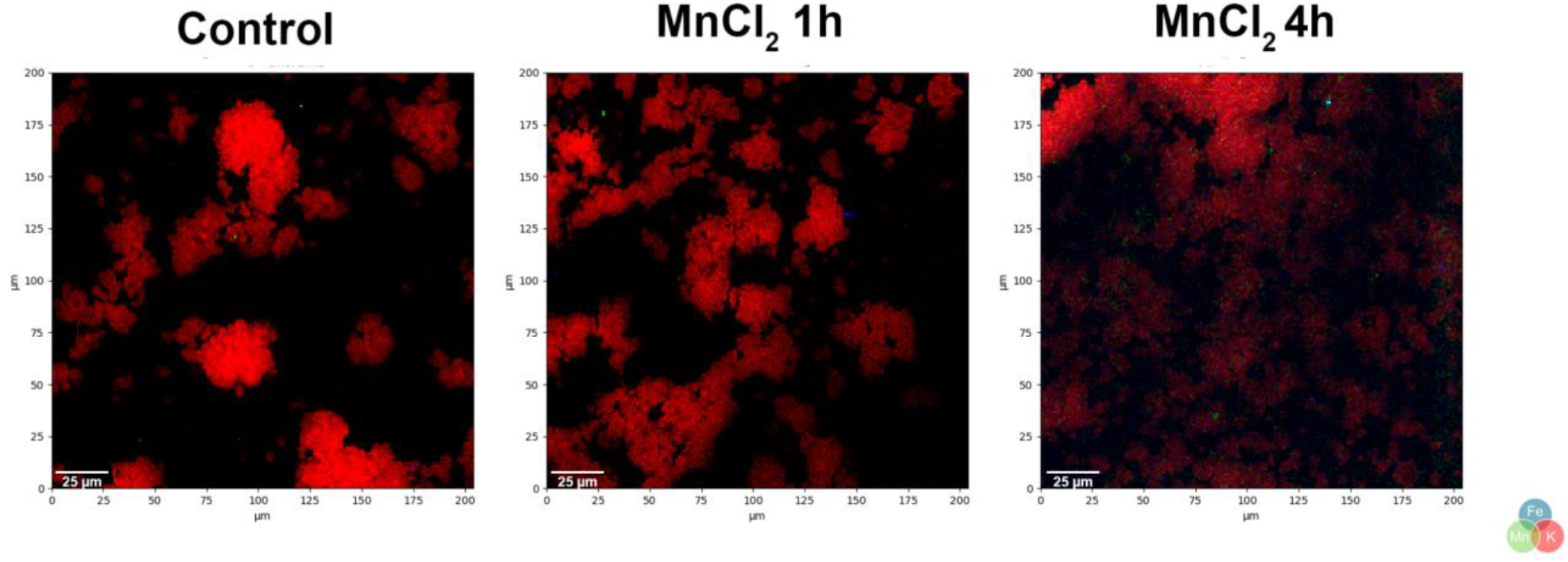
Panorama multielemental maps of LLC cells exposed to control media or supplemented with MnCl_2_ for 1 or 4h. Scale bar = 25 µm. Color code: Potassium (K): red, Iron (Fe): blue, and Manganese (Mn): red.

Interestingly, the distribution of Mn in the cell monolayer of the high Mn group occurs heterogeneously, similar to previous data obtained from resected tumors^33^. Next, we decided to investigate the niches of interest using the flyscan mode. In this mode, the maps are generated in higher resolution (100 nm), covering a 100 × 100 µm area. Briefly, in this mode, Mn accumulation becomes more evident (Figure 5).

**Figure 5.**
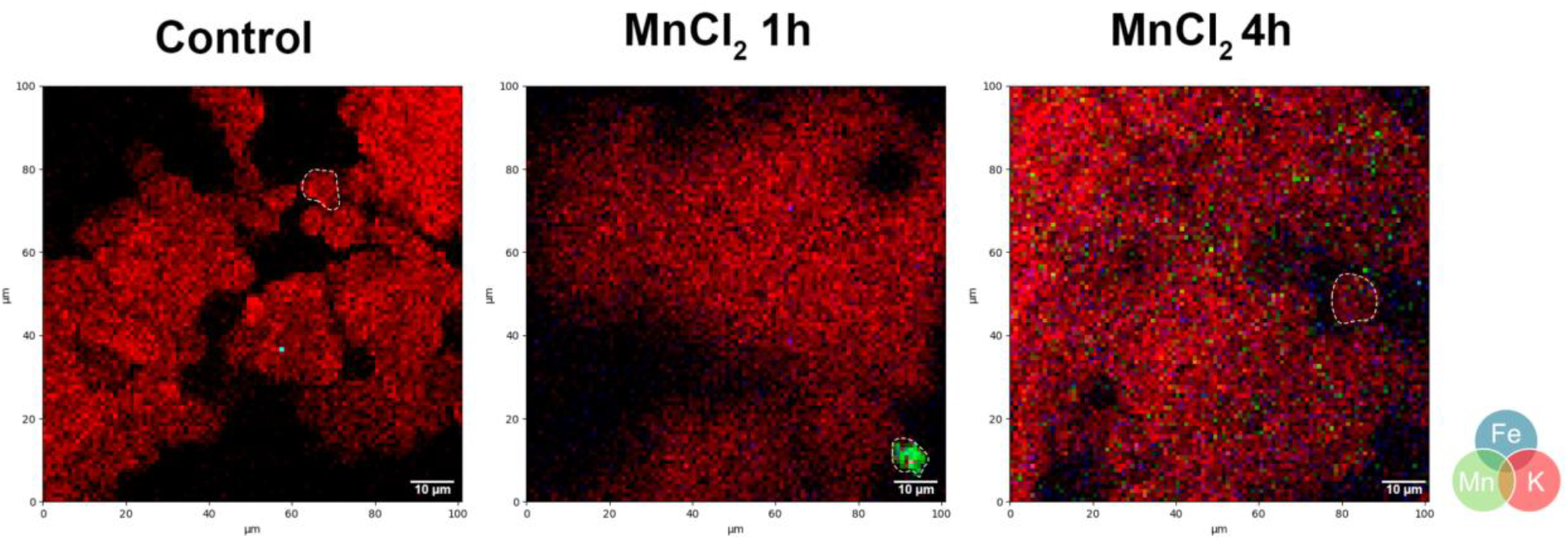
Representative flyscan multielemental maps of LLC cells exposed to control media or supplemented with MnCl_2_ for 1 or 4h. Scale bar = 10 µm. Dashed lines indicate individual cells. Color code: Potassium (K): red, Iron (Fe): blue, and Manganese (Mn): red.

For the control group, all regions irradiated in flyscan mode presented a wide distribution of cells, highlighted by the element K. As expected, we observed a few niches rich in Mn and Fe, with superposition with the K signal, suggesting a natural accumulation of these elements in cells (Figure 6). Cells exposed to MnCl_2_ for 1h presented an increase in the number and intensity of Mn and Fe niches overlapping the K signal (Figure 6). While the 4h group demonstrated a higher distribution of Mn niches for the samples, suggesting that Mn interacts with cells in a time-dependent manner (Figure 6).

**Figure 6.**
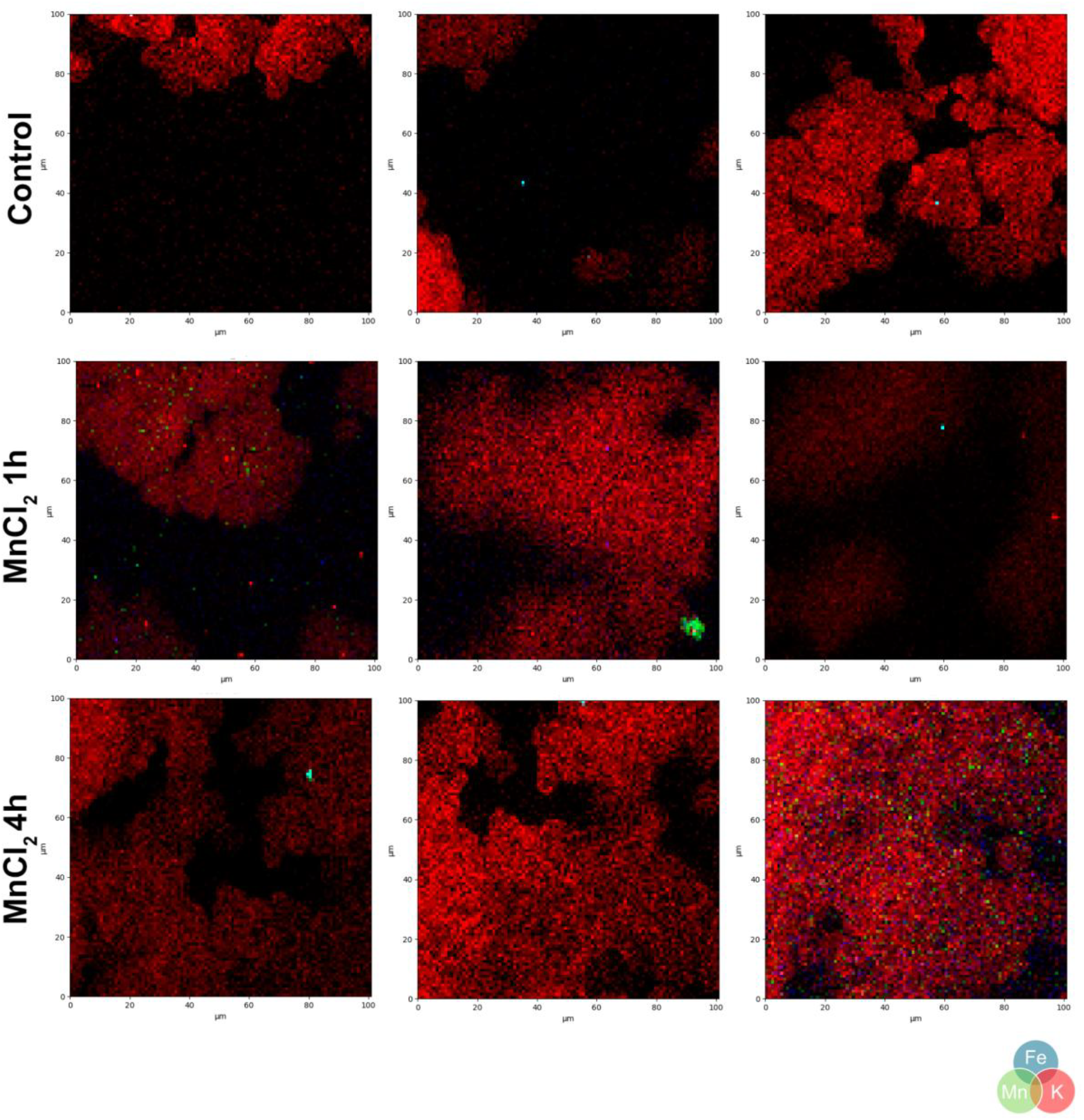
Representative flyscan multielemental maps of LLC cells exposed to control media or supplemented with MnCl_2_ for 1 or 4h. Color code: Potassium (K): red, Iron (Fe): blue, and Manganese (Mn): red.

Recent evidence suggests a new feature of tumors, such as the accumulation of Mn and Fe in elevated but non-toxic concentrations. We previously observed that LLC cells pre-exposed to Mn are capable of retaining the element and secreting it on extracellular vesicles^33^.

Now, we demonstrate that these cells are capable of rapid internalization of divalent cations, specifically Mn (Table 1 and Figure 3). In about 1h, cells had 11X more Mn internalized than their respective control, as shown in Figure 3, A. Mn accumulation and tumor malignancy were associated previously with the generation of metastatic niches in vivo ^33,55^, chronic exposure was related to mammary gland hyperplasia in mice^56^, but also with radioresistance in glioblastoma and melanoma human samples^17^. It was also reported that increased Mn levels were found in tumor tissue from colorectal cancer patients compared to adjacent non-tumoral tissue^57^.

Mechanistically, Mn accumulation relies on Mn transporters in both plasmatic and organelle membranes, such as SLC11A2, SLC39A8, and SLC39A14^58–61^, for example. These transmembrane proteins regulate Mn homeostasis by controlling Mn levels in organelles, the extracellular environment, and the cytoplasm. One potential hypothesis is that tumoral Mn-accumulative tissues present elevated expression of these transmembrane proteins. SLC11A2 gene codifies for the transmembrane protein Divalent Metal Transporter 1 (DMT1), found on the cell membrane and endosomes^62^, is mostly known for Fe transport, but DMT1 has also affinity for Mn and Cu ions^63^, with expression regulated by Mn availability according to the work of Bai and collaborators^64^. SLC39A8 and SLC39A14 encode ZIP8 and ZIP14 (Zrt, Irt-like protein 8 and 14), respectively, which are mostly known for Zn and Fe transport^58,65,66^. However, both transporters were described to regulate Mn content in mammalian cells, as reviewed by Winslow *et al*. in 2020 and Fujoshiro & Kambe in 2022^59,67^. Like DMT1, ZIP8 and 14 expression are also responsive to Mn exposure in cellular and murine models^68,69^, reinforcing our hypothesis of increased Mn levels related to transporter expression on cell surface. Indeed, the expression of both transporters in the blood-brain barrier is associated with Mn accumulation in the brain and neurotoxicity^70^.

DMT1 mRNA is found overexpressed in colorectal cancer samples from TCGA, and is related to increased Fe uptake in mice. DMT1 selective disruption in this model led to decreased Fe uptake, reduced tumor growth, and tumor cell proliferation^71^. Pan-cancer analysis also identified DMT1 overexpression in ovarian cancer samples and further validated with LC-MS, ELISA, and immunohistochemistry in patient samples; gene knockout reduced and inhibited tumor cell proliferation and migration, respectively^72^. ZIP8 mRNA is upregulated in 35 of 40 cancers analysed in TCGA database, including lung adenocarcinoma^73^. *In vitro* evaluation of ZIP8 protein levels in healthy and tumoral lung lineages indicated that ZIP8 overexpression is pertinent for tumoral cells^73^. Knockdown of ZIP8 induced cell cycle arrest in neuroblastoma SH-SY5Y lineage and modulated cell proliferation negatively, associated with reduced nuclear translocation of NF-kB^74^. Immunohistochemistry analysis of HCC samples showed an increase in ZIP14 abundance and correlated with poor prognosis^75^. Interestingly, p53 suppression was correlated with ZIP14 overexpression and increased levels of Fe uptake^76^. The authors discuss the relevance of this event since many tumors are characterized by p53 loss and how tumor metabolism could be deeply altered in this condition.

Despite the evidence presented about the importance of these transporters in cancer, some of these works did not perform a multielemental quantification of metals. In this case, tumor malignancy relies only on transporter expression, but not on the metals transported by them. Our data show that, for lung adenocarcinoma model, only Mn is accumulated in cells (Figure 3, A), and as evidenced by XRF results, the population of Mn-cumulative cells is heterogeneous, suggesting a subpopulation of cells with potential to become more malignant, as previously observed by the group. Future studies in our group aim to investigate Mn intracellular route, as well as transporter expression, especially in the Mn-cumulative subpopulation.

### 3. Cells exposed to high-Mn conditions present major migratory activity

To understand the effect of these elements on cell behavior, we performed a wound healing assay. We evaluated cells’ ability to migrate while cultivated in medium supplemented with 5 µM of each salt. Corroborating with previous data, cells grown in MnCl_2_ medium had the highest percentage of relative distance migrated (35.9%) (Figure 7). While cells cultivated with CuCl_2_ (20.6%), FeCl_2_ (30.1%), and ZnCl_2_ (26.1%) did not present significant changes in migration compared to the control group (23.7%) (Figure 7).

**Figure 7.**
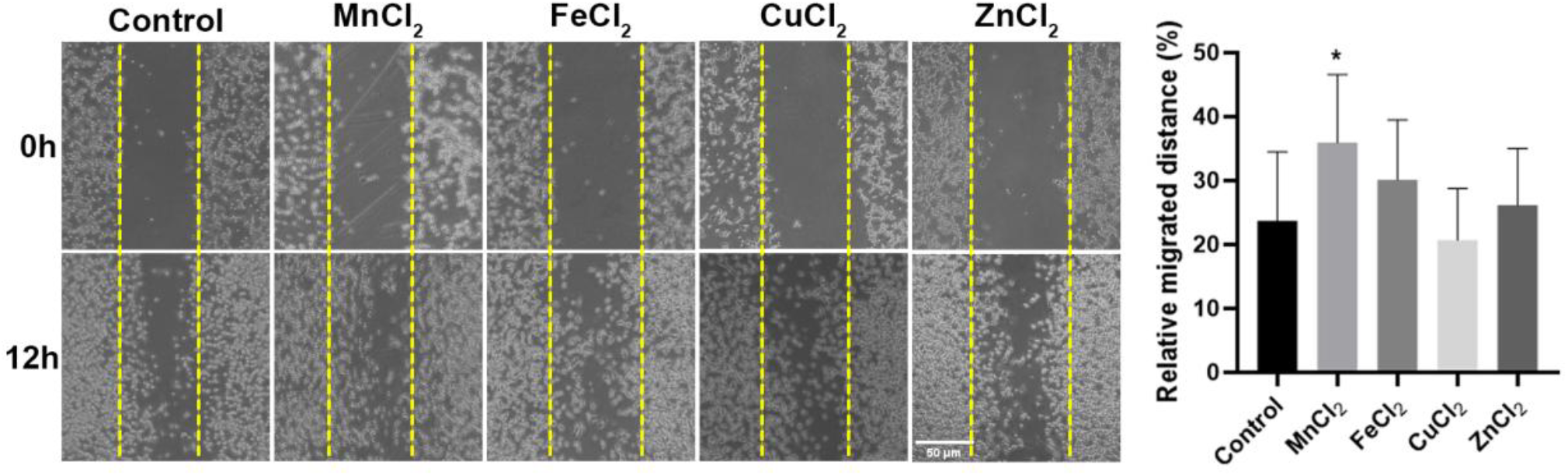
Relative migrated distance after 12h exposure to control media or supplemented with MnCl_2_, FeCl_2_, CuCl_2_, or ZnCl_2_. Scale bar = 50µm. *p≤0,05. Mean±standard-deviation. n=9. One-Way ANOVA and Dunnett post-test.

Wound healing assay results reflect the importance of metals to cell adhesion and migration. Divalent cations are potent cofactors to regulate integrin activity^77^. Cells exposed to Mn presented 35.9% of relative migration, 12% higher than the control group, suggesting a major coactivation of integrins, as suggested before by the group when an increased colocalization rate between β1-integrin and syndecan-1^33^ was observed. Mn^2+^ ions increase the affinity of integrins with their ligands^78^, being fundamental for integrin α4β1 and chondroitin sulfate proteoglycans^79^. Mn^2+^ interaction with integrins was proved to enhance tumor malignancy in the models of melanoma, glioblastoma, and breast cancer through increasing adhesion^80–82^ and migration^83^. In our model, cells exposed to Fe and Cu did not present increased migration. However, Fe ions were supposed to regulate tumor cell invasiveness via intracellular pathways, as observed in breast cancer cells^84,85^, as well as leading to EMT in the models of breast, lungs, and pancreatic cancer^85–87^. Although Cu is related to malignancy, previous data suggest that the element is involved in tumor angiogenesis and TME remodeling, as in collagen synthesis^88,89^ or through growth factor signalling^90,91^. From the metals tested in this work, Zn was the one with relative migration most similar to the control group (26.1% and 23.7%, respectively), as expected. According to the work of Lymburner, Zn^2+^ ions are potent inhibitors of cell migration and adhesion, regulating negatively integrin activation, in an opposite role as proposed for Mn^2+^ and Mg^2+^ ions^83^. Zn role in malignancy is not related to invasion, but for regulating essential processes for tumor sustained proliferation, such as gene transcription, DNA synthesis, and apoptosis regulation^92–94^. Zn dyshomeostasis in tumors is frequently associated with mutations or altered expression in Zn transporters^95–97^, which are associated with other TME changes that contribute to increased proliferation, differentiation, and migration^5,98–100^.

## Conclusion

In this work, we aimed to understand the effect of tumor metallome modulation. We characterized the resistance of LLC cells to toxic concentrations of Fe, Zn and Cu, while maintaining sensitivity to Mn toxicity. We also evaluated metal internalization, and distribution as well as cell migration. The results presented highlight the influence of Mn initial interactions with tumor cells, reinforcing previous data obtained by our group. LLC cells quickly accumulate and respond to the increased Mn availability, resulting in a malignant phenotype.

## Methods

### 1. Cell culture

LLC (mouse Lewis lung carcinoma, ATCC - CRL-164) cells were cultivated in standard conditions, 37ºC and 5% CO_2_ atmosphere. Cell medium contained DMEM (Dulbecco’s Modified Eagle medium - Vitrocell) supplemented with fetal bovine serum (FBS) 10% (v/v) (Vitrocell) and glucose (ISOFAR) 4.5 g/L. Cells were passed with trypsin solution (Sigma-Aldrich) 0.5%.

### 2. Cell viability assay (MTT)

Cells were seeded in a 96-well plate (2.0×10^3^ / well). After reaching 85% confluence, cells were cultivated in standard medium (control) or supplemented with 1.5, 3.1, 6.2, 12.5, 25, 50, 100, or 500 µM of MnCl_2_, FeCl_2_, CuCl_2_, or ZnCl_2_. After 24h, cells were incubated with MTT (3-(4,5-dimethyl-2-thiazolyl)-2,5-diphenyl-2H-tetrazolium bromide) reagent (Invitrogen M6494) 0.5 mg/mL in cell medium for 1h. Next, the medium was discarded, and formazan crystals were dissolved in 100 µL dimethyl sulfoxide (DMSO, Sigma-Aldrich) per well. Absorbance at 520 nm was acquired using SpectraMax Plus microplate reader (Molecular Devices) and analyzed using Softmax Pro software (Molecular Devices). Results were expressed in viability percentage compared to the control group.

### 3. Clonogenic assay

LLC cells were seeded in 35 mm plates (5,0×10^2^/plate) in control medium or supplemented with 1, 5, 50, or 500 µM of MnCl_2_, FeCl_2_, CuCl_2_, or ZnCl_2_. 24h later, the cell medium was exchanged for regular medium, and the growth of colonies was monitored for 7 days. Next, cells were fixed in absolute ethanol and stained with crystal violet (Merck 111885/1) 1:15 (v/v) in 100% ethanol. The colonies were registered and counted using ImageJ software.

### 4. Metal content quantification through Inductively Coupled Plasma Optical Emission Spectrometry (ICP-OES)

Cells were seeded in 35 mm plates (1.0×10^5^/plate) and grown until they reached 95% confluence. Then, they were exposed to control medium or supplemented with 5 µM of MnCl_2_, FeCl_2_, CuCl_2_, or ZnCl_2_ for 1h or 4h. 1 mL. Next, 1 mL of cell supernatant was collected, and cells were washed with PBS (phosphate saline buffer) at room temperature and lysed with H_2_O.

The cell lysate was centrifuged for 15 min at 3500 RPM, and the supernatant was collected for ICP-OES analysis. All samples were analyzed in Perkin Elmer Optima 7300 DV ICP-OES. The following parameters were used for the analyses: peristaltic pump flow 1.5 mL/min, plasma flow 15 L/min, nebulization flow 0.6 L/min, 1400 W, radial plasma view for Ca and K, axial plasma view for Cu, Fe, Mn, P, and Zn. Multielemental calibration curve ranged from 0.001 to 20.0 mg/L.

### 5. High-resolution X-ray fluorescence (XRF) analysis

XRF analysis was performed at Sirius Light Source at the Brazilian Synchrotron Light Source Laboratory (LNLS). Cells were seeded in Si_3_N_4_ bases (Norcada NX5200D) until complete adhesion and then exposed to control medium or supplemented with MnCl_2_ 5 µM for 1h or 4h. Next, the medium was aspirated, cells were washed once with PBS at room temperature, and air-dried. Samples were irradiated at the Tarumã substation of Carnaúba (Coherent X-Ray Nanoprobe Beamline) beamline at room temperature, ambient pressure, and kEV. Multielemental maps were produced with PyMca software (version 5.6.5).

### 6. Cell migration wound healing assay

Cells were seeded in 35 mm plates (1,5×10^5^/plate). When cells reached 90% confluence, the medium was removed, and cells were mechanically removed from the plate in a cross-shaped pattern using a sterile P1000 tip. Next, cells were carefully rinsed to remove debris and incubated with control medium or supplemented with 5 µM of MnCl_2_, FeCl_2_, CuCl_2_, or ZnCl_2._ To evaluate cell migration and wound closure, cells were imaged, using the cross-center as a reference, at 0 h and 12 h. Migration was quantified using ImageJ software (version 1.74K) and expressed as the percentage of migrated distance relative to the original wound width.

### 7. Statistical analyses

Statistical analyses were performed with GraphPad Prism 8 for Windows, version 8.0.2 (2019). One-way ANOVA test with Tukey post-test or Sidak’s multiple comparison post-test was applied where applicable.

